# Ecology and transmission of a dengue virus serotype 4 identified in wild *Aedes aegypti* in Florida

**DOI:** 10.1101/2021.07.11.451177

**Authors:** Jasmine B. Ayers, Xuping Xie, Heather Coatsworth, Caroline J. Stephenson, Christy M. Waits, Pei-Yong Shi, Rhoel R. Dinglasan

## Abstract

1

Dengue virus is the most prevalent mosquito-borne virus, causing approximately 390 million infections and 25,000 deaths per year. *Aedes aegypti*, the primary mosquito vector of dengue virus, is well established throughout the state of Florida, USA. Autochthonous transmission of dengue virus to humans in Florida has been increasing since 2009, alongside consistent importation of dengue cases. However, most cases of first infection with dengue are asymptomatic and the virus can be maintained in mosquito populations, complicating surveillance and leading to an underestimation of disease risk. Metagenomic sequencing of *Aedes aegypti* mosquitoes in Manatee County, Florida revealed the presence of dengue virus serotype 4 (DENV-4) genomes in mosquitoes from multiple trapping sites over 2 years, in the absence of a human DENV-4 index case and even though a locally acquired case of DENV-4 has never been reported in Florida. This finding suggested that: i) DENV-4 may circulate amongst humans undetected, ii) the virus was being maintained in the mosquito population, or iii) the detected complete genome sequence may not represent a viable virus. This study demonstrates that an infectious clone generated from the Manatee County DENV-4 (DENV-4M) sequence is capable of infecting mammalian and insect tissue culture systems, as well as adult female *Aedes aegypti* mosquitoes when fed in a blood meal. However, the virus is subject to a dose dependent infection barrier in mosquitoes, and has a kinetic delay compared to a phylogenetically related wild-type (WT) control virus from a symptomatic child, DENV-4H (strain *Homo sapiens*/Haiti-0075/2015, GenBank accession MK514144.1). DENV-4M disseminates from the midgut to the ovary and saliva at 14 days post-infection. Viral RNA was also detectable in the adult female offspring of DENV-4M infected mosquitoes. These results demonstrate that the virus is capable of infecting vector mosquitoes, is transmissible by bite, and is vertically transmitted, indicating a mechanism for maintenance in the environment without human-mosquito transmission. These findings suggest undetected human-mosquito transmission and/or long-term maintenance of the virus in the mosquito population is occurring in Florida, and underscore the importance of proactive surveillance for viruses in mosquitoes.

**Graphical Abstract:** In order to better assess the public health risk posed by a detection of DENV-4 RNA in Manatee County, FL *Aedes aegypti*, we produced an infectious clone using the sequence from the wild-caught mosquitoes and characterized it via laboratory infections of mosquitoes and mosquito tissues.

## 2 Introduction

Dengue virus (DENV) is a single stranded, positive-sense RNA arthropod-borne virus in the flavivirus family, which is predominantly transmitted in human populations by the bite of infected female *Aedes aegypti* mosquitoes. DENV causes dengue fever, the most common arboviral disease in humans with approximately 40% of the world’s population at risk of infection (Scientific Working Group on Dengue. Meeting (: Geneva and UNDP/World Bank/WHO Special Programme for Research and Training in Tropical Diseases, 2007). Four antigenically distinct serotypes of DENV circulate in human populations (denoted DENV-1 through DENV-4). Primary DENV infections are typically asymptomatic or cause non-descript, febrile illness, but subsequent re-infection with a different serotype can cause severe dengue, including dengue shock syndrome, dengue hemorrhagic fever, or death (Guzman and Vazquez, 2010). *Aedes aegypti* and a secondary dengue vector *Aedes albopictus* are widespread in Florida, USA (Reiskind and Lounibos, 2013). Imported (i.e., travel-related) cases of dengue occur in Florida every year, and locally acquired cases have been increasing over the past decade with 73 locally acquired cases in the state in 2020 alone (Mosquito-Borne Disease Surveillance | Florida Department of Health). Travel associated cases of all 4 serotypes of DENV have been reported in Florida, but no locally acquired cases of DENV-4 have ever been reported (Mosquito-Borne Disease Surveillance | Florida Department of Health).

In 2016 and 2017, we detected and subsequently sequenced the complete genome of dengue virus serotype 4 (DENV-4) from pools of field-derived F1 *Aedes aegypti* mosquitoes collected in Manatee County, Florida (Boyles et al., 2020). The presence of DENV-4 in mosquitoes in two consecutive years from the same oviposition traps without a locally acquired or travel associated human index case in the county was peculiar, suggesting either undetected human-to-mosquito transmission and/or prolonged maintenance of the virus via vertical transmission from mosquitoes to their progeny. Natural vertical transmission in *Aedes* has been reported for all 4 DENV serotypes, although its contribution to epidemic disease in humans is unclear (Ferreira-de-Lima and Lima-Camara, 2018). However, the discovery of DENV-4 from Manatee County field-derived mosquitoes was unexpected since the project was focused on characterizing the adult female *Ae. aegypti* microbiome and metavirome, targeting insect-specific RNA viruses in a county assumed to be well-outside the DENV infection foci of South Florida. The mosquito samples were processed solely for RNA extraction, library assembly and subsequent sequencing, and as such, arbovirus isolation was not possible due to viral inactivation during the sample preparation process.

We generated an infectious clone virus to complete the rigorous validation of the reported viral genome sequenced from the Manatee *Ae. aegypti* using validated methods to produce flavivirus infectious clones (Shan et al., 2016; Shi et al., 2002; Zou et al., 2015; Xia et al., 2018). The clone was based on the published virus genome (Accession Number: MN192436.1) to characterize the viability of the resulting virus DENV-4M. Herein, we demonstrate that the DENV-4M infectious clone is indeed a viable virus, which is infectious to both susceptible mammalian and mosquito cell lines. Importantly, we observed that the infectious clone can infect *Ae. aegypti* Orlando strain mosquitoes *per os* and is secreted into saliva at 14 days post-infection. The virus also infects the ovary 14 days following *per os* infection and can undergo vertical and transstadial transmission. This suggests that the initial identification of DENV-4M genomes in Manatee County mosquitoes represented a real detection of DENV-4, which changes our understanding of the public health risk of this serotype in the state. Given the possible risk of transmission to humans by bite as well as long-term maintenance in the mosquito population by vertical transmission, these data further underscore the importance of pro-active surveillance for DENV and other arboviruses in vector mosquito populations in advance of the “mosquito season” in Florida.

## 3 Materials and Methods

### 3.1 Construction of DENV-4M infectious clone

Four fragments (FI-FIV) spanning the entire genome of DENV-4M (GenBank accession:MN192436.1) were initially synthesized and cloned into pUC57 vector by Genscript (Piscataway, NJ). A T7 promoter and a hepatitis delta virus ribozyme (HDVr) sequence were engineered at the 5′ and 3′ ends of the fragment FI and FIV respectively. The individual fragment was amplified by PCR using the Platinum SuperFi II DNA Polymerase (ThermoFisher Scientific, Waltham, MA) with corresponding primer pairs listed in Table 1. The resulting amplicons were assembled into a full-length clone in a single-copy vector pCC1BAC (Epicentre) by using the NEBuilder HiFi DNA Assembly kit (New England Biolabs, Ipswich, MA). The cDNA sequence of DENV-4M in the full-length clone was finally validated by Sanger sequencing using the primers listed in Table S2.

### 3.2 RNA *in vitro* transcription, electroporation and virus rescue

Full-length DENV-4M RNAs were *in vitro* transcribed using a T7 mMessage mMachine kit (ThermoFisher Scientific, Waltham, MA) from cDNA plasmids as linearized by ClaI. The RNA transcripts (10 μg) were electroporated into BHK-21 cells following a protocol described previously (Xie et al., 2011) with some modifications. Briefly, 8×10^6^ cells were suspended in 800 μl Ingenio® Electroporation Solution (Mirus Bio, Madison, WI) and mixed with 10 μg RNA in a 4-mm cuvette. Electroporation was performed using the GenePulser apparatus (Bio-Rad) by three pulses with 3 s intervals at instrumental settings of 0.85 kV at 25 μF. After a 10-min recovery at room temperature, transfected cells were transferred into a T-75 flask containing 15 ml culture media. Alternatively, 1×10^4^ transfected cells were seeded into each well of 8-well Lab-Tek™ II chamber slides (ThermoFisher Scientific) for immunostaining analysis. After incubation at 37°C with 5% CO_2_ for 24 h, the culture medium was replenished with medium contanining 2% FBS. The cells were then incubated at 30°C with 5% CO_2_ for additional 4 days. Supernatants were clarified by centrifuging at 1000 g for 5 min at 4°C and stored at −80°C prior to use.

### 3.3 Cell and virus culture

BHK-21 cells (Xie et al., 2011) and Vero E6 cells (ATCC# C1008) were obtained directly from Pei-Yong Shi’s group at UTMB. Vero E6 cells were maintained at 37°C and 5% CO2 in complete DMEM (ThermoFisher, 11965092) supplemented with 10% heat inactivated FBS, 1x penicillin/streptomycin (ThermoFisher, 15140122) and 1x L-glutamine (ThermoFisher, 25030081). Aedes albopictus C6/36 (ATCC# CRL-1660) and *Ae. aegypti* Aag2 (ATCC# CCL-125) insect cell lines were obtained from ATCC and maintained at 30°C and 5% CO2 in complete MEM (ThermoFisher, 12360-038) supplemented with 10% heat inactivated FBS, 1x penicillin/streptomycin and 1x L-glutamine. The DENV-4M was obtained from UTMB as passage 0 infectious virus at 4 × 10^3^ PFU/mL in Vero E6 cell culture supernatant and was used directly in experiments as described. A dengue virus serotype 4 strain isolated from a symptomatic child in Haiti in 2015 (DENV-4H, strain Homo sapiens/Haiti-0075/2015, GenBank accession MK514144.1) was used as a positive control.

### 3.4 *In vitro* infection

Cells were infected at 80% confluency at 0.01 multiplicity of infection in media (DMEM or MEM as described in cell and virus culture section) containing 3% FBS (i.e., reduced serum media). After a 1-hour infection period, inoculum was removed, the monolayer was washed once with 1x PBS, and fresh reduced serum media was added. For rt-qPCR experiments, supernatant samples were taken at this point (0dpi). Mock infected controls were seeded and treated identically, with sterile media used instead of virus stock. To passage virus, 200 μL of 5dpi culture supernatant from the previous passage’s infected flask was used as the virus stock for the new flask. Supernatant samples were collected during passages 1, 2, 5, and 10. All cell culture experiments were performed in triplicate and each independent replicate was a culture slide well or flask processed in parallel.

### 3.5 Mosquito rearing and *in vivo* infection

ORL strain mosquitoes were initially obtained as adults from the Gainesville United States Department of Agriculture Center for Medical, Agricultural, and Veterinary Entomology colony. Offspring of these mosquitoes were used for experiments. Adults were maintained on 10% sucrose solution ad libitum at 28°C and 80% humidity with a 12:12 light:dark cycle. Larvae were reared on ground TetraMin flakes. Adult female mosquitoes were starved overnight and fed a 2:2:1 mixture of O+ human red blood cells (Lifesouth Community Blood Centers, Gainesville, FL): infected or uninfected (in naive blood control conditions) Vero E6 cell culture supernatant: heat inactivated human serum via an artificial membrane feeder held at 37°C and affixed with pork sausage casing. After blood feeding, mosquitoes were cold anesthetized and non-blood engorged individuals were discarded. All remaining mosquitoes were maintained on 10% sucrose solution and given access to an oviposition surface. At 7 and 14dpi, mosquitoes were cold anaesthetized, surface sterilized in 70% ethanol, and rinsed twice in 1x PBS before midguts and ovaries were dissected out in 1x PBS. In the vertical transmission experiment, mosquitoes were offered a second naive blood meal at 13dpi, and allowed to oviposit again for 72h on new damp filter paper. These second blood feed eggs were dried completely, hatched, and reared to adulthood (as described above). Resultant adult females were surface sterilized and pooled by rearing container in pools of up to 25 (Table S1). Each replicate (2 or 3 replicates as indicated in figure legends) used mosquitoes from a different egg laying date, which were reared, fed, and processed separately. All mosquito infections and handling of infected mosquitoes took place in an ACL-3/BSL-2 facility.

### 3.6 Salivation assay

At 14dpi, DENV-4 infected mosquitoes were starved overnight. To collect saliva, mosquitoes were cold-anesthetized, and their wings and legs were removed. Each mosquito was then fastened to a glass microscope slide with tape, and their proboscis was inserted into a graduated glass capillary tube (Drummond, Broomall, PA) filled with 3μL of warmed human O+ blood (1:1 O+ human red blood cells: heat inactivated human serum) to initiate feeding cues and facilitate saliva collection (Stephenson et al., 2021). The mosquitoes were placed in a lit rearing chamber at 28°C with 80% relative humidity for forty-five minutes or until they ingested approximately 2μL of blood. Each proboscis was then removed from its capillary tube, and the remaining blood from each capillary tube was aspirated into 1.5mL microcentrifuge tubes with 200μL of reduced (3% FBS) DMEM. Mosquitoes were then surface sterilized in 70% ethanol and rinsed twice in 1x PBS before being dissected in PBS to produce paired ovary and midgut samples. All samples were immediately stored at −80°C until use.

### 3.7 RNA extraction and reverse transcription quantitative PCR (rt-qPCR) virus detection

At the time of collection, each tissue was placed in a 1.5mL microcentrifuge tube with 700μL chilled, sterile PBS and 0.2mL of sterile glass beads. Each tissue sample was loaded into a Bullet Blender and homogenized by running the Bullet Blender at speed 8 for 5 minutes. Saliva, supernatant, and tissue samples were then spun down in a bench-top centrifuge at 3750xg for 3 minutes. Lysis buffer (560μL) AVL (Qiagen) was aliquoted into pre-labelled sterile 1.5mL microcentrifuge tubes. 140μL of sample homogenate was aliquoted into the corresponding lysis tube. Cell supernatant samples were added directly into lysis buffer. RNA extraction on each sample was carried out using the QIAmp Viral RNA extraction kit (Qiagen) following the manufacturer’s protocols. Sample RNA was tested for dengue virus serotype 4 (DENV-4) pre-membrane protein gene **(primer and probe sequences are provided in Table S3)**. Each sample was run as technical duplicates, and each plate included a no template control, and a DENV-4 positive control (NR-50533, BEI resources and diluted 1:10 with nuclease-free water). Sample RNA was run with either QuantaBio UltraPlex 1-Step ToughMix (4X) Low-ROX master mix, or SuperScript™ III Platinum™ One-Step qRT-PCR on a BioRad CFX96 Touch Real-Time PCR Detection System at 50°C for 30 minutes (for Superscript reactions) or 50°C for 10 minutes (QuantaBio reactions), 95°C for 2 minutes, and 45 cycles of: 95°C for 15 seconds, and 60°C for 45 seconds.

The rt-qPCR CT values were converted to PFU equivalents (PFUe) with a standard curve created using eight ten-fold dilutions of RNA extracted from DENV-4M Vero E6 P2 stock virus of known titer (7 × 10^6^ PFU/mL), run as technical duplicates, and fitted with a logarithmic line of best fit in Microsoft Excel (Version 2105) ((*CT value*) = −1.542ln(*PFUe*) + 39.355, R² = 0.9905). The limit of detection (LOD) for this assay was a CT value of 40, or 0.65 PFUe/mL.

### 3.8 Immunofluorescence assays

Unless otherwise indicated, all incubation steps were performed at 4°C in the dark, and all buffers were kept ice cold. Media was washed off of 4dpi cells, and 7dpi or 14dpi midguts or ovaries with 1x PBS. Tissues were fixed by adding 1mL 4% paraformaldehyde (PFA) in PBS + 0.05% Tween20 (PBST) and leaving cells at room temperature for 10 minutes. PFA/PBST mixture was removed, and tissue was washed 3 × 5 minutes with 1mL of PBST. Tissues were permeabilized by adding 1mL 0.5% TritonX-100/PBST for 20 minutes. TritonX-100/PBST mixture was removed and tissue was washed 3 × 5 minutes with 1mL of PBST. Tissues were blocked in 1 mL of 5% heat-inactivated fetal bovine serum (FBS) in PBST for 30 minutes at room temperature. FBS/PBST mixture was removed and tissues were washed 3 × 5 minutes with 1mL PBST. Primary antibody (200 μL) in PBS was added (1:2,000 dilution of pan-serotype DENV NS1 mAb (R&D Biosystems# MAB94442-100) in 1x PBS) and tissues were incubated in a humidity chamber overnight at 4°C. Primary antibody was removed and tissues were washed 3 × 5 minutes with 1mL of PBST. Secondary antibody (200 μL) was added (1:1,000 dilution of Alexa Fluor™ goat anti-mouse 594 IgG (H+L) (Invitrogen, A11005, Lot 1937185) in 1x PBS) and tissues were incubated for 1 hour at 4°C in the dark. Cells were washed 3 × 5 minutes with 1mL of PBST. DAPI stain was added (1:200 dilution of Roche Diagnostics, Ref 10236276001, Lot 70317525 in 1x PBS) and tissues were incubated for 10 minutes at room temperature in the dark. Tissues were rinsed 2x with PBST and washed once for 5 minutes with PBS. Slides were mounted with VECTASHIELD® Antifade Mounting Media with DAPI (Vector Laboratories, Ref H-1200, Lot ZE0815) and imaged on a KEYENCE BZ-X800 microscope. Capture and image processing settings were kept constant between conditions within each timepoint/tissue.

### 3.9 Plaque assay

Supernatant samples used as virus stocks to infect cells or mosquitoes were subject to titration by plaque assay and were mixed with 10% final concentration trehalose to stabilize virions for freezing and stored in liquid nitrogen. Baby hamster kidney fibroblast (BHK-21) cells were grown to confluency with DMEM supplemented with 10% FBS, 1% L-glutamine, 1x penicillin/streptomycin and 0.25 μg/mL amphotericin B and then seeded into 24-well plates and incubated for two days at 37°C and 5% CO2. Next, each supernatant sample was serially diluted 10-fold in reduced DMEM (3% FBS). The spent media was removed from the BHK-21 cells in the 24-well plates and 100μL of each dilution series was added to individual wells. The 24-well plates were then rocked at room temperature for 15 minutes and incubated at 37°C and 5% CO2 for 45 minutes. Afterwards, 500μL of 0.8% w/v methyl cellulose in DMEM containing 2% FBS was added, and the plates were re-incubated at 37°C and 5% CO2 for five days. On the fifth day, the spent media was removed from each well of the 24-well plates and a 1:1 methanol/acetone solution with 1% crystal violet was added for at least one hour to fix and stain the cells. The plates were then washed with water, and subsequently stored upside down overnight to drain and dry them. Plaques were manually counted, and titer expressed as plaque forming units/mL (PFU/mL).

## 4 Results

### 4.1 Construction of the Manatee DENV-4 Infectious Clone (DENV-4M)

Construction of full-length infectious cDNA clones of flaviviruses remains challenging due to the instability of the viral genome during plasmid propagation in the *Escherichia coli* (*E.coli*) system. We took two steps to overcome this issue. Firstly, to quickly obtain the subclones prior to assembly of the full-length infectious clone, we divided the entire DENV-4M cDNA into four consecutive fragments and cloned them into a high-copy plasmid pUC57 (Fig. 1A). To enable the *in vitro* transcription of a 5′ capped genome-length RNA, a T7 promoter and a hepatitis delta virus ribozyme (HDVr) sequence were engineered upstream of the 5′ untranslated region (UTR) and downstream of the 3′ UTR, respectively. Secondly, upon assembly we used high-fidelity PCR to obtain each fragment, and took advantage of the NEBuilder HiFi DNA Assembly technique to clone the four PCR amplicons into a single-copy vector pCC1BAC to increase the stability of the cDNA plasmids when propagated in *E. coli*. Nineteen-to 26-basepair (bp) overlaps were introduced into adjacent fragments. In addition, the 5′ end of PCR fragment FI and the 3′ end of PCR fragment FIV contain a 24-bp overlap with the region upstream of the restriction site *NotI* and a 21-bp overlap with the region downstream of the restriction site *ClaI* in the pCC1BAC vector, respectively (Fig. 1A). The four fragments were then directionally assembled into the pCC1BAC that was pre-linearized by NotI and ClaI, resulting in the full-length infectious clone pCC1-DENV-4M FL.

**Figure 1:**
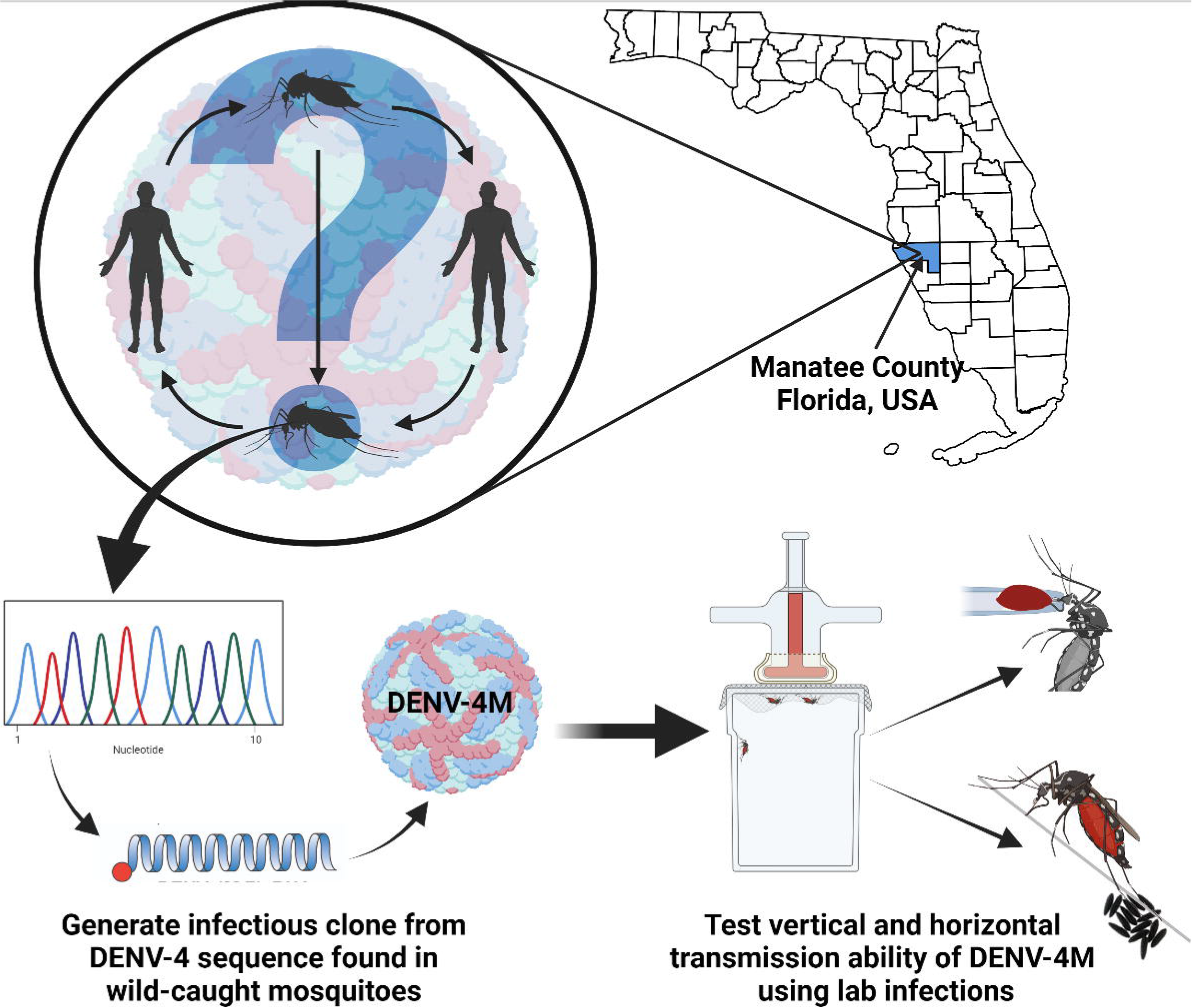
DENV-4M parental infection data shows active replication in mammalian cell culture. **(a)** Diagram of construction of DENV-4M infectious clone and generation of recombinant viruses. **(b)** IFA analysis of BHK-21 cells transfected with *in vitro* transcribed viral RNA. On day 1, 3, and 5 after transfection, cells were assayed by immunofluorescence for DENV nonstructural protein 4B (red). Nuclei are stained with DAPI (blue).

To recover recombinant DENV-4M from the infectious clone pCC1-DENV-4M FL, we electroporated the *in vitro* transcribed genome-length RNA into BHK-21 cells. After electroporation, the intracellular expression of nonstructural protein 4b (NS4B) was examined by immunofluorescence assay (IFA). NS4B-positive cells increased from day 1 to 5 post-electroporation (Fig. 1B). These data demonstrated that DENV-4M is rescued from the infectious clone and the resulting recombinant DENV-4M virus can replicate and spread on BHK-21 cells. The virus was expanded once on Vero E6 cells and then utilized for the experiments described in the remainder of the study.

### 4.2 *In vitro* immunofluorescence assay shows DENV-4M replicates in mammalian and insect cell lines

To assess the viability of DENV-4M in insect tissues *in vitro*, an *Aedes albopictus* embryonic cell line known to be highly permissive to DENV-4 infection (C6/36) was chosen as a model insect line. African green monkey kidney cells (Vero E6) were used as a positive control. IFA was performed for DENV-4 nonstructural protein 1 (NS1) to visualize viral replication in cell cultures. DENV-4M produced NS1 signal by 4 days post infection (dpi) in both C6/36 **(Fig. 2a-b)** and Vero E6 cells **(Fig. 2c-d)**. A DENV-4 strain isolated in 2015 from a symptomatic child in Haiti (DENV-4H, strain *Homo sapiens*/Haiti-0075/2015, GenBank accession MK514144.1) was used throughout the study as a positive infection control, as it is known to infect *Ae. aegypti* robustly (Stephenson et al., 2021). In both cell lines, DENV-4M showed noticeably lower infection prevalence and intensity compared to DENV-4H **(Fig. 2)**.

**Figure 2:**
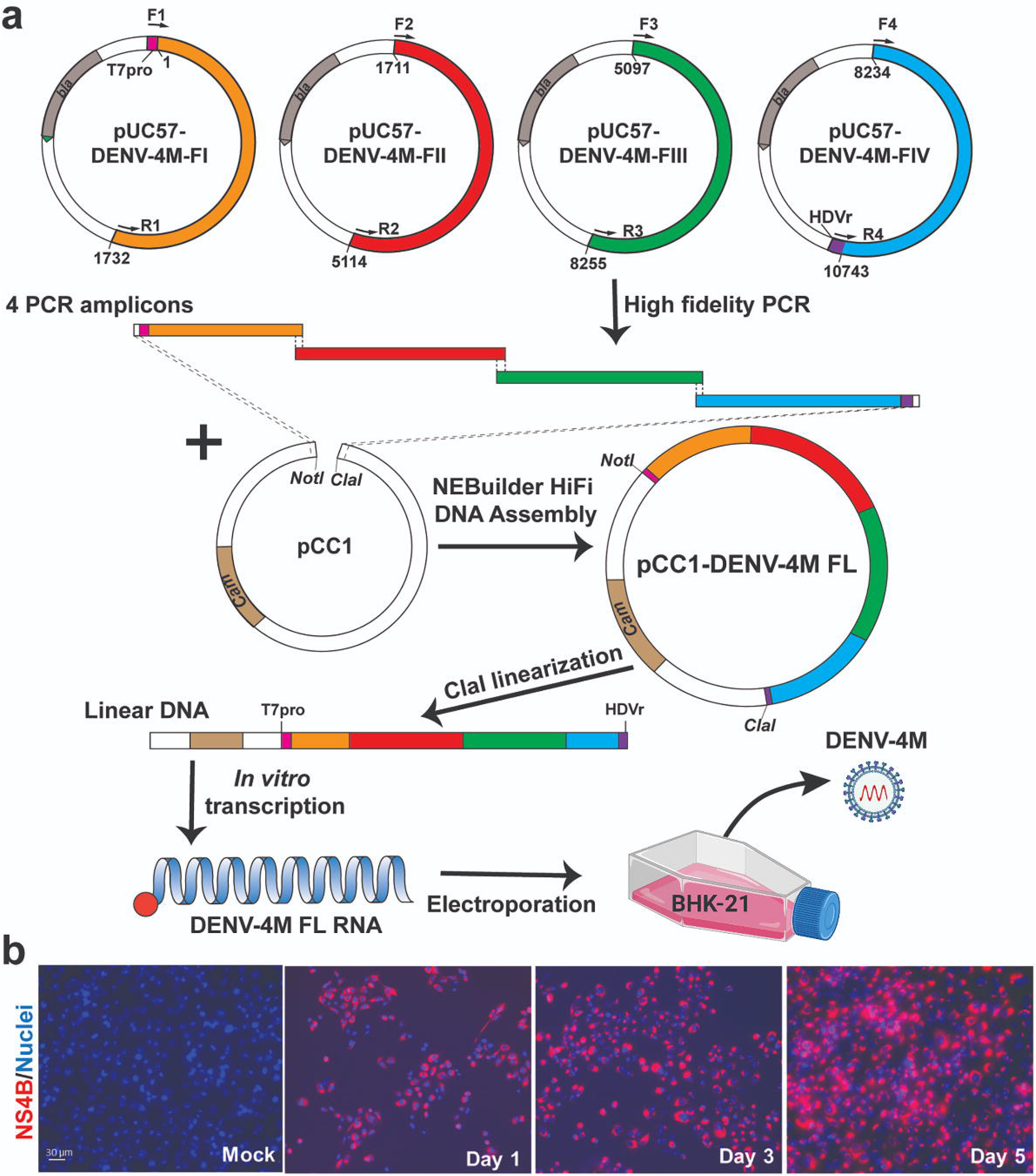
DENV-4 NS1 protein IFA shows viral replication in *Ae. albopictus* C6/36 cells and in mammalian Vero E6 cells. DENV-4M or DENV-4H were used to infect C6/36 **(a-b)** and Vero E6 cells **(c-d)** at an MOI of 0.01 and cells were processed for NS1 IFA at 4dpi. Red (center column) indicates DENV-4 NS1 and blue (left column) indicates DAPI DNA counterstain. Representative images were chosen from 3 independent experiments. Scale bar = 100μm.

### 4.3 Replication rate in insect cell lines can be improved by serial passage

To quantify the replication rate of DENV-4M *in vitro*, we performed reverse transcription quantitative PCR (rt-qPCR) on RNA from culture supernatant collected immediately after inoculum was washed off cells (0dpi), and on supernatant collected 5 days post-infection (5dpi). To confirm that viable virus was being produced, we passaged supernatant from passage 1 (P1) flasks into fresh cultures and repeated the rt-qPCR quantification **(Fig. 3a)**. By this measure, DENV-4M replicated robustly in Vero E6 cells with an average 10^3.95^ (8,947-fold) increase in PFU equivalents (PFUe)/mL from 0dpi to 5dpi in P2 cultures **(Fig. 3b)**. However, replication in C6/36 cells was much more modest with a maximum PFUe/mL increase of 10^1.95^ (89-fold) in the P2 cultures, with one of three replicates not producing viable progeny virus in P1 **(Fig. 3b)**. A third cell line, the *Ae. aegypti* larval cell line Aag2, did not demonstrate any replication in P1 or P2 **(Fig. 3b)**. Despite being a more relevant model (in that they are derived from *Ae. aegypti*), Aag2 cells were expected to be less permissive to replication than C6/36 cells because they have an intact RNA interference pathway while C6/36 cells do not (Brackney et al., 2010).

**Figure 3:**
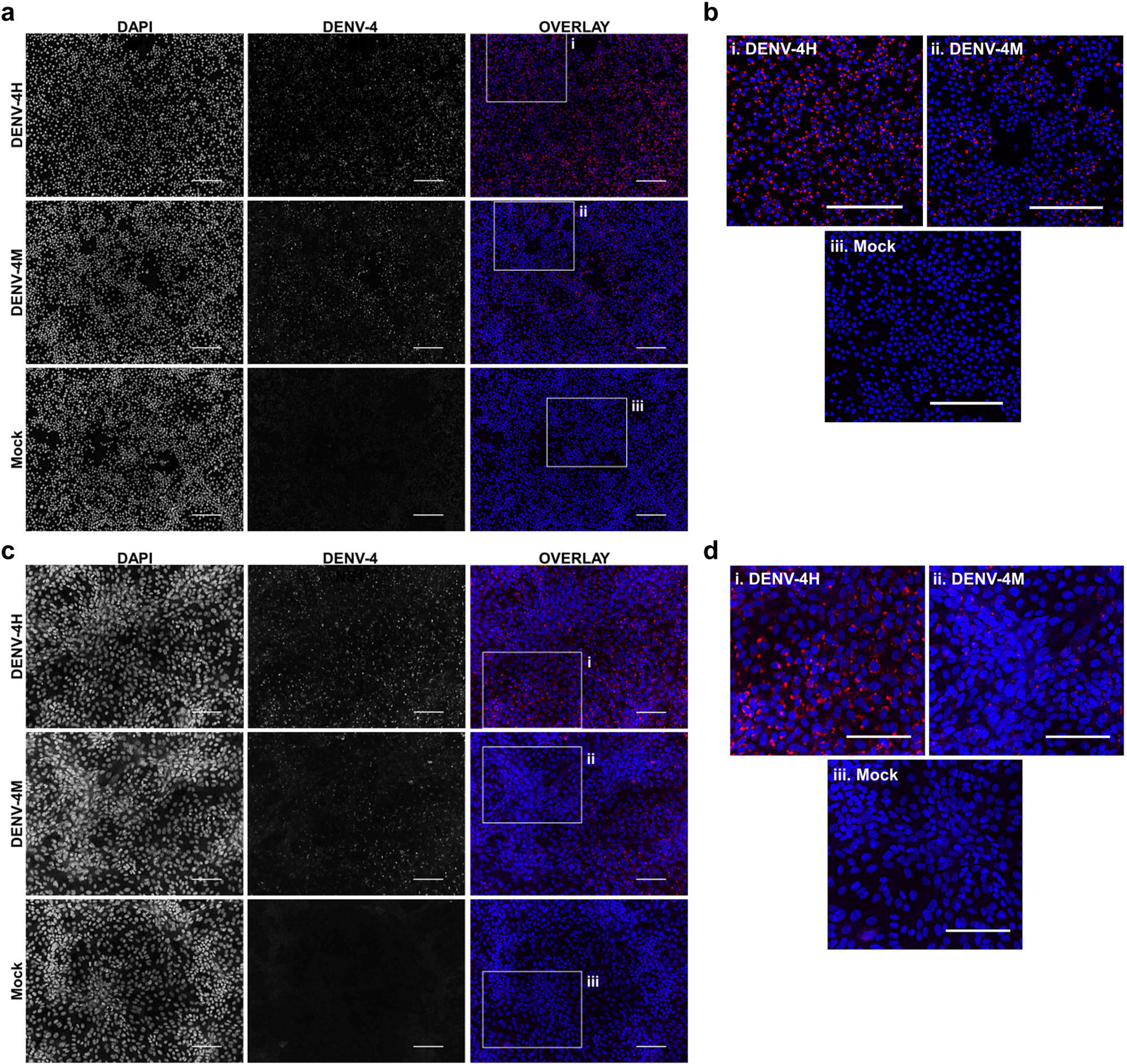
Detection of Manatee IC replication in mammalian and insect cells by rt-qPCR on culture supernatant shows DENV-4M completes replication in insect cell lines and can be adapted to insect cell culture by serial passage. **(a)** Workflow schematic of serial passage experiment. One mammalian (Vero E6) and 2 mosquito (C6/36 and Aag2) cell lines were infected with parental (P0) DENV-4M stock virus. Cell culture supernatant was taken at day 0 (immediately following removal of inoculum) and day 5 post infection for rt-qPCR. On day 5 post infection, supernatant from the infected culture was serially passaged onto an uninfected culture. Additionally, C6/36 P5 virus was used to infect Aag2 cell culture to test if adaptation to an insect cell line improves the performance of the virus in Aag2 cells. **(b)** Log_10_ DENV-4 PFU equivalents (PFUe)/mL, as determined by converting cycle threshold values with a standard curve made using 10-fold dilutions of RNA extracted from virus stock with a known PFU of 7 × 10^6^ are reported for the indicated conditions. The values reported represent the mean of 2 technical rt-qPCR replicates. ND indicates that no viral genome was detected in the indicated condition. The limit of detection of the assay was a cycle threshold value of 40, or 0.65 PFUe/mL The three biological replicates reported were performed in separate tissue culture flasks processed in parallel.

As the virus replicated more robustly in P2 than P1 on C6/36 cells in two of three replicates, we continued to serially passage infected supernatant on C6/36 cells to adapt the virus to insect cell culture. At P5 and P10, samples of day 0 and day 5 supernatant were retained for rt-qPCR analysis; DENV-4M replicated much more rapidly in C6/36 cells by P10 with a maximum PFUe/mL increase of 10^3.13^ (1,363-fold), comparable to its initial replication rate in Vero E6 cells **(Fig. 3b)**. DENV-4M transferred to Aag2 cells after P5 on C6/36 cells also fared better than the P0 parental stock, with two of three replicates showing replication and a maximum PFUe/mL increase of 10^1.51^ (32-fold). The ability of DENV-4M to establish replication in insect cells over serial passage was sporadic compared to Vero E6 cells, with one of three replicates in both C6/36 and Aag2 cells not demonstrating replication in cell cultures after P1 **(Fig 3b)**.

### 4.4 DENV-4M (Vero E6 P2) is detectable by rt-qPCR *in vivo* and is capable of horizontal and vertical transmission

To obtain an *in vivo* measure of viral replication, we initially fed an infectious blood meal containing parental P0 DENV-4M to adult female Orlando strain (ORL) mosquitoes and dissected midguts at 14 dpi, however, no midguts were found to be positive (0/43) for DENV-4 by this measure.

Based on the weak but detectable replication observed in the initial infection of C6/36 cells and the low titer of parental DENV-4M (4 × 10^3^ PFU/mL), we hypothesized that the resistance to infection observed in ORL mosquitoes may be dose-dependent. To test this, we fed adult female ORL mosquitoes an infectious blood meal containing DENV-4M from P2 on Vero E6 cells (7 x10^6^ PFU/mL) (experimental workflow illustrated in **Fig. 4a**). DENV-4M is clearly capable of replicating in ORL mosquitoes following 2 passages on Vero E6 cells **(Fig. 4b)**. Midguts were collected from these mosquitoes as a proxy for infection, saliva was collected as a proxy for horizontal transmission, and ovaries were used as a proxy for vertical transmission. Infected mosquitoes showed a high infection intensity and prevalence in the midgut and ovary on day 14 **(Fig. 4b)**. Of the tested mosquitoes, 19/68 (27.9%) had detectable viral genomes in the saliva at day 14, suggesting that the potential for horizontal transmission of DENV-4M by bite exists **(Fig. 4b)**. Based on evidence that multiple blood feeding increases virus dissemination in *Ae. aegypti* (Armstrong et al., 2020), we provided a cohort of these mosquitoes a second non-infectious blood meal at 4dpi, but this did not improve infection prevalence or intensity in this model **(Fig. S1)**.

**Figure 4:**
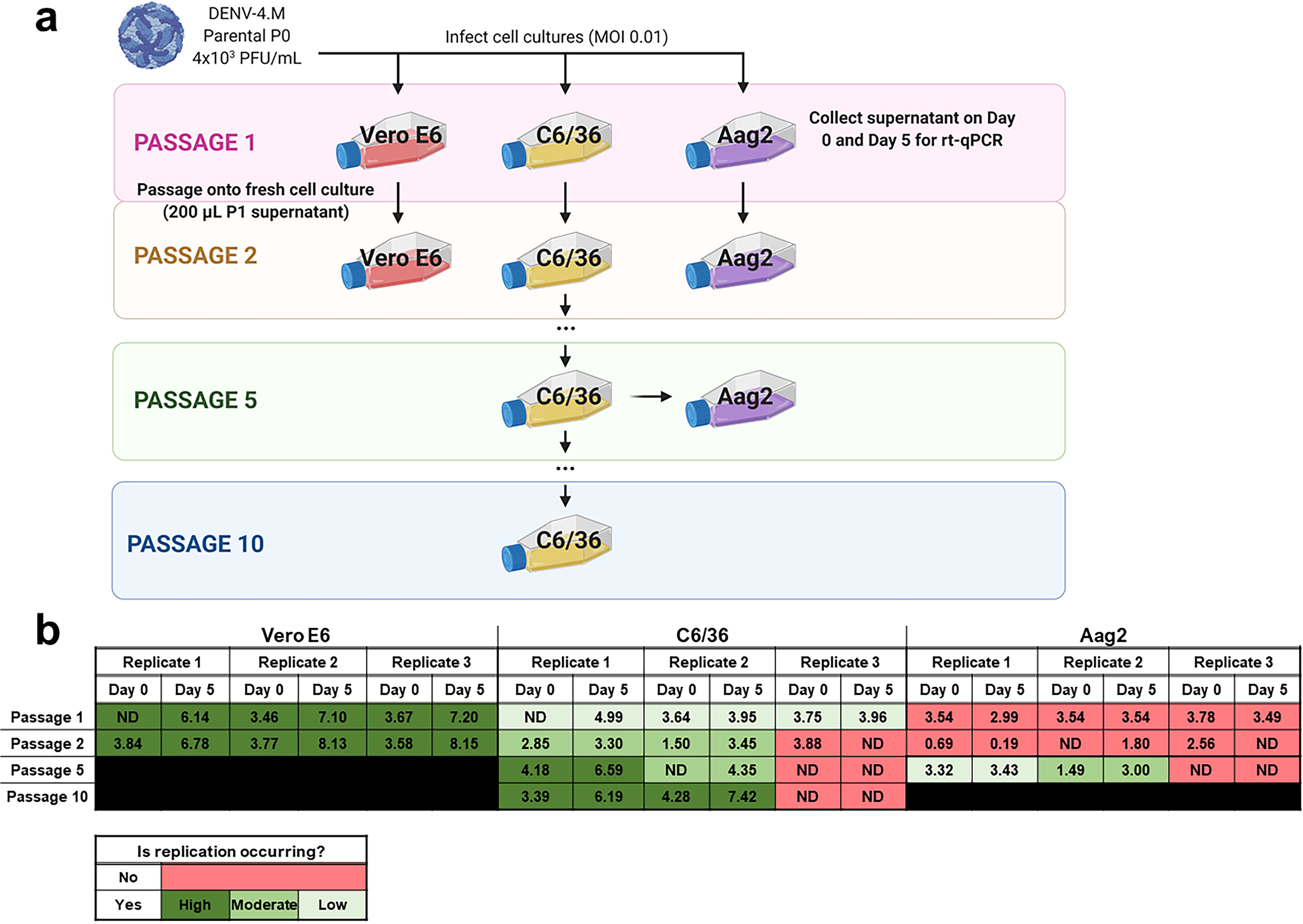
Detection of DENV-4M genome by rt-qPCR after *per os* infection in 14dpi ORL mosquito tissues. **(a)** Workflow of *in vivo* infection. Parental P0 DENV-4M stock yielded no infection in 43 tested 14dpi midguts. Virus was passaged 2x in Vero E6 cells and ORL mosquitoes were infected with this product. **(b)** Rt-qPCR was performed on saliva, ovaries, and midguts dissected from each of 68 individual mosquitoes. Log_10_ DENV-4 PFU equivalents (PFUe)/tissue, as determined by converting cycle threshold (CT) values with a standard curve made using 10-fold dilutions of RNA extracted from virus stock with a known PFU/mL of 7 × 10^6^ are reported. The values reported represent the mean of 2 technical rt-qPCR replicates. The limit of detection (LOD) of the assay was a CT value of 40, or 0.65 PFUe/tissue. The three biological replicates reported were performed in separate tissue culture flasks processed in parallel. Only samples with a detectable CT value in both technical replicates are shown on the graph; the proportion of rt-qPCR positive samples / total samples tested is displayed over each tissue type. Results are pooled from 3 independent experiments (saliva, ovaries, and midgut) or 2 independent experiments (adult female progeny).

To test whether vertical transmission was possible, we provided ORL females fed Vero E6 P2 DENV-4M a non-infectious blood meal at 13 dpi to initiate a second gonotrophic cycle, collected and hatched eggs, and reared the resulting F1 progeny to adulthood. Twelve pools of up to 25 surface-sterilized adult female F1s **(Table S1)** were used for virus detection by rt-qPCR. Of these, only 1 of the 12 tested pools was positive for DENV-4, indicating that DENV-4M can undergo vertical transmission in *Ae. aegypti* infected *per os* **(Fig. 4b)**, but efficiency is low.

### 4.5 DENV-4M replicates in the midgut and ovaries of adult female mosquitoes after an infectious blood meal

To confirm the dissemination of DENV-4M to the ovary deduced from the rt-qPCR data, as well as to compare the kinetics of this process to the wild type virus DENV-4H, adult female ORL mosquitoes were fed blood meals containing DENV-4 H, Parental P0 DENV-4M, and Vero E6 P2 DENV-4M at high titer (7 × 10^6^ PFU/mL) or low titer equivalent to that of the parental virus (4 × 10^3^ PFU/mL). At 7 and 14dpi, midguts and ovaries were dissected from these mosquitoes, and DENV-4 replication was visualized using a DENV NS1 IFA. In both organ types, trachea displayed red autofluorescence as can be seen in the naive blood fed control panels. Image capture and analysis settings were held constant across conditions within each tissue and timepoint. At 7dpi, only DENV-4H infection was visible in midguts **(Fig. 5 a-b)** and ovaries **(Fig. 5 d-e)**. No discernable NS1 signal was seen in DENV-4M infected mosquitoes at the 7dpi timepoint. The major structures of a DENV-4 negative ovariole are indicated in **Fig. 5c**. At 14 dpi DENV-4M high titer and DENV-4H produced a strong NS1 stain in the midgut **(Fig. 6 a-b)**. DENV-4M high titer produced an NS1 stain in the secondary follicles of the ovaries at 14 dpi while viral replication in the ovaries of DENV-4H mosquitoes was largely abated by this timepoint **(Fig. 6 c-d)**. Neither the parental P0 or the Vero E6 P2 low titer DENV-4M stocks produced noticeable NS1 stain in the ovaries or midgut at either timepoint, confirming that DENV-4M is not capable of infecting mosquitoes *per os* at the low titer, and that the infectivity difference between DENV-4M parental P0 and DENV-4M Vero E6 P2 is likely due to the different titer rather than genotypic differences between the virus stocks which arose during passage of the virus on Vero E6 cells **(Fig. 5–6)**.

**Figure 5:**
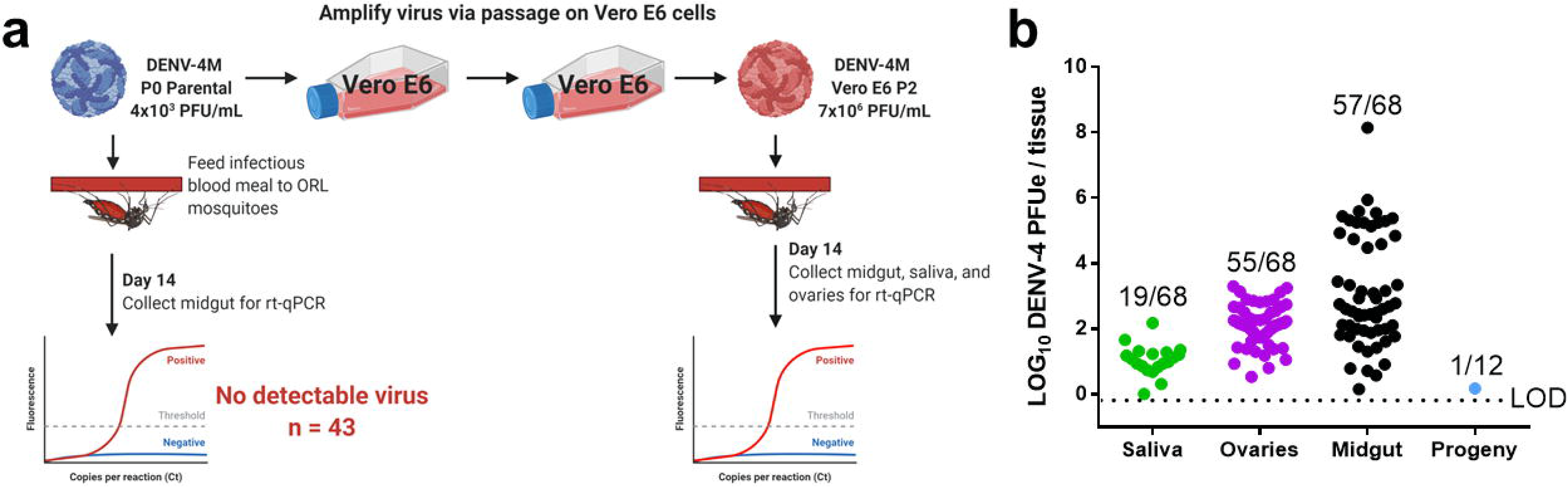
DENV-4H replicates in mosquito midguts and ovaries at 7dpi, but establishment of DENV-4M infection is delayed. Adult female ORL mosquitoes were fed a blood meal containing DENV-4H (5 × 10^6^ PFU/mL), Parental P0 DENV-4.M (3 × 10^4^ PFU/mL), Vero E6 P2 DENV-4M (high titer: 7 × 10^6^ PFU/mL, low titer: 4 × 10^3^ PFU/mL), or naive blood without virus. On day 7 post-infection, midguts **(a-b)** and ovaries **(d-e)** were dissected, and virus replication was visualized by NS1 IFA (red, middle column) with DAPI DNA counterstain (blue, left column). **(c)** Shows the major structures of a DENV-4 negative ovariole in DAPI/NS1 IFA and bright field; the primary follicle/developing embryo, the undeveloped secondary follicle containing nurse cells, and the germarium. **(b)** and **(e)** are insets chosen to show NS1 signal in virus positive conditions, compared to the naive blood negative control. Representative images were chosen from at least 2 independent replicates. Scale bar = 100μm.

**Figure 6:**
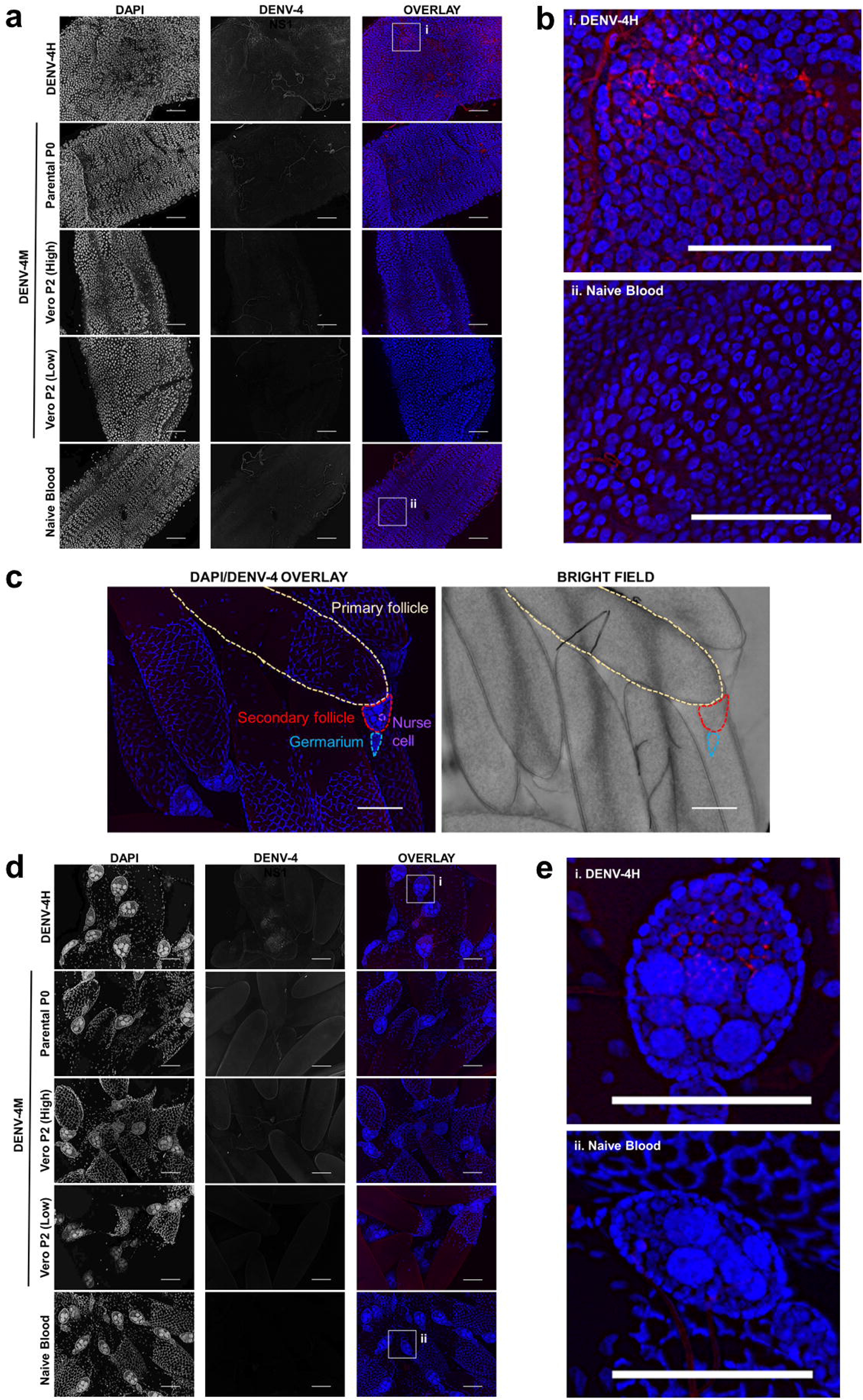
DENV-4M replicates in the midgut and ovary at 14dpi. Adult female ORL mosquitoes were fed a blood meal containing DENV-4H (5 × 10^6^ PFU/mL), Parental P0 DENV-4M (3 × 10^4^ PFU/mL), Vero E6 P2 DENV-4M (high titer: 7 × 10^6^ PFU/mL, low titer: 4 × 10^3^ PFU/mL), or naive blood without virus. On day 14 post-infection, midguts **(a-b)** and ovaries **(c-d)** were dissected, and virus replication was visualized by NS1 IFA (red, middle column) with DAPI DNA counterstain (blue, left column). **(b)** and **(d)** are insets chosen to show the NS1 signal in virus positive conditions, compared to the naive blood negative control. Representative images were chosen from at least 2 independent replicates. Scale bar = 100μm.

## 5 Discussion

To gain a better estimate of mosquito transmission potential of the Manatee County, FL *Aedes aegypti* DENV-4 in the absence of directly isolated virus, we characterized an infectious clone generated from the *Ae. aegypti* field-derived arboviral genome sequence. Gaining retrospective insight into arbovirus transmission dynamics and disease risk in a local setting is especially important considering that the original DENV-4M virus was detected from *Ae. aegypti* collected from a tourist corridor in Manatee County across two consecutive years and in the absence of DENV-4 human index cases (Boyles et al., 2020; Mosquito-Borne Disease Surveillance | Florida Department of Health).

The data presented here indicate that DENV-4M is a viable virus capable of replicating in insect and mammalian cell lines as well as in adult female Orlando strain *Aedes aegypti* mosquitoes after *per os* infection. DENV-4M is present in saliva in this model, posing a risk of transmission to humans by bite. It also disseminates to the ovary and is detectable in infected F1 adult female progeny, indicating vertical and transstadial transmission, which could facilitate long term maintenance within Manatee *Ae. aegypti*.

DENV-4M replication kinetics show slower and less robust replication than the wild type DENV-4H control. There was a clear dose dependent infection barrier for DENV-4M in the mosquito model as neither the parental P0 or Vero E6 P2 virus replicated in the examined tissues with the low 4 × 10^3^ PFU/mL titer. However, the high titer used herein is within the range of blood titers seen in viremic humans (Xu et al., 2020), so this dose-dependent infection barrier does not preclude mosquitoes from acquiring DENV-4M from infected human hosts.

Since the viral genome sequence was from nulliparous mosquitoes reared from oviposition traps and ostensibly must have undergone vertical transmission, our expectation was that it would be well adapted to replicating in insect tissue. Therefore, DENV-4M’s strong *in vitro* preference for Vero E6 cells over insect cells was surprising. However, since the virus was originally propagated in Vero E6 cells, an initial sub-selection for variants performing well in this mammalian cell line may have been inadvertently performed. This hypothesis is supported by the observation that the virus’ replication rate in insect cell lines could be markedly improved by serial passage. The sporadic nature of this adaptation to insect cell lines is illustrated by the fitness differences in virus stocks from replicate experiments, suggesting that the genetic bottleneck produced by serial passage influences the virus’ performance by inducing genetic drift, as has been previously observed in other RNA viruses (Duarte et al., 1992; Chao, 1990; Clarke et al., 1993; Lázaro et al., 2003). These results imply that minority variants, which arose randomly during passage of DENV-4M, vastly altered the virus stock’s phenotype even in only one or two passages. Genetic drift is perhaps of particular import in arboviruses since their transmission cycle requires them to maintain infectiousness in two extremely divergent hosts, and this host switching has been observed to constrain their genetic diversity (Moutailler et al., 2011; Coffey and Vignuzzi, 2011; Grubaugh et al., 2016). A comparison of the genome sequences of virus lineages derived from a single known parental genome sequence, which either succeeded or failed to adapt to the infection of mosquito cells may be a useful approach for identifying virulence factors that mediate virus infectivity for *Ae. aegypti* and is a compelling topic for future study of DENV-4M. The data suggest that generating infectious clones for use in arbovirus vector competence studies in insect cell lines rather than mammalian cell lines should be considered, as this may avoid a reduction in virus fitness in mosquitoes.

This study demonstrated that infectious clones are extremely useful research tools, particularly to test viability when a viral genome is obtained but live virus isolation was not possible. However, when characterizing infectious clone-derived viruses, care should be taken to assess how the severe genetic bottleneck produced first by reducing the original virus population to a single consensus sequence, as well as during the cell culture process, may affect the phenotype. Taken together, these results demonstrate that our identification and sequencing of DENV-4M from mosquitoes from a central-southwest Florida county represents a bona fide maintenance of the virus in this environment, compelling a recalibration of the perceived risk of DENV transmission in the state.

## 7 Author Contributions

X.X. designed and constructed the DENV-4M infectious clone and generated parental DENV-4M virus stocks. J.B.A. designed and performed mosquito and cell line infection experiments as well as the bulk of sample processing, imaging, and data analysis. X.X. and J.B.A. generated figures. H.C. and C.J.S. performed salivation assays. H.C. also helped perform preliminary experiments to optimize and validate IFA and imaging methods. C.M.W. contributed to processing RNA samples. P.S. and R.R.D. provided guidance on project design and writing. All authors contributed to the writing of the manuscript, but it was predominantly written by J.B.A. and X.X. with significant editing contributions from H.C. and R.R.D.

## 9 Acknowledgments

DENV-4 Haiti (DENV-4H) stocks were generously provided by John Lednicky’s lab at the University of Florida. Baby hamster kidney fibroblast (BHK-21) cells used for plaque assays were a kind gift from the Dimopoulos Lab at Johns Hopkins. The DENV-4 positive control was obtained through BEI Resources, NIAID, NIH: Genomic RNA from Dengue Virus Type 4, UIS 497, NR-50533. Figures were constructed using Biorender.com and GraphPad Prism 6 (graphs).

---

**Supplementary Figure 1:**
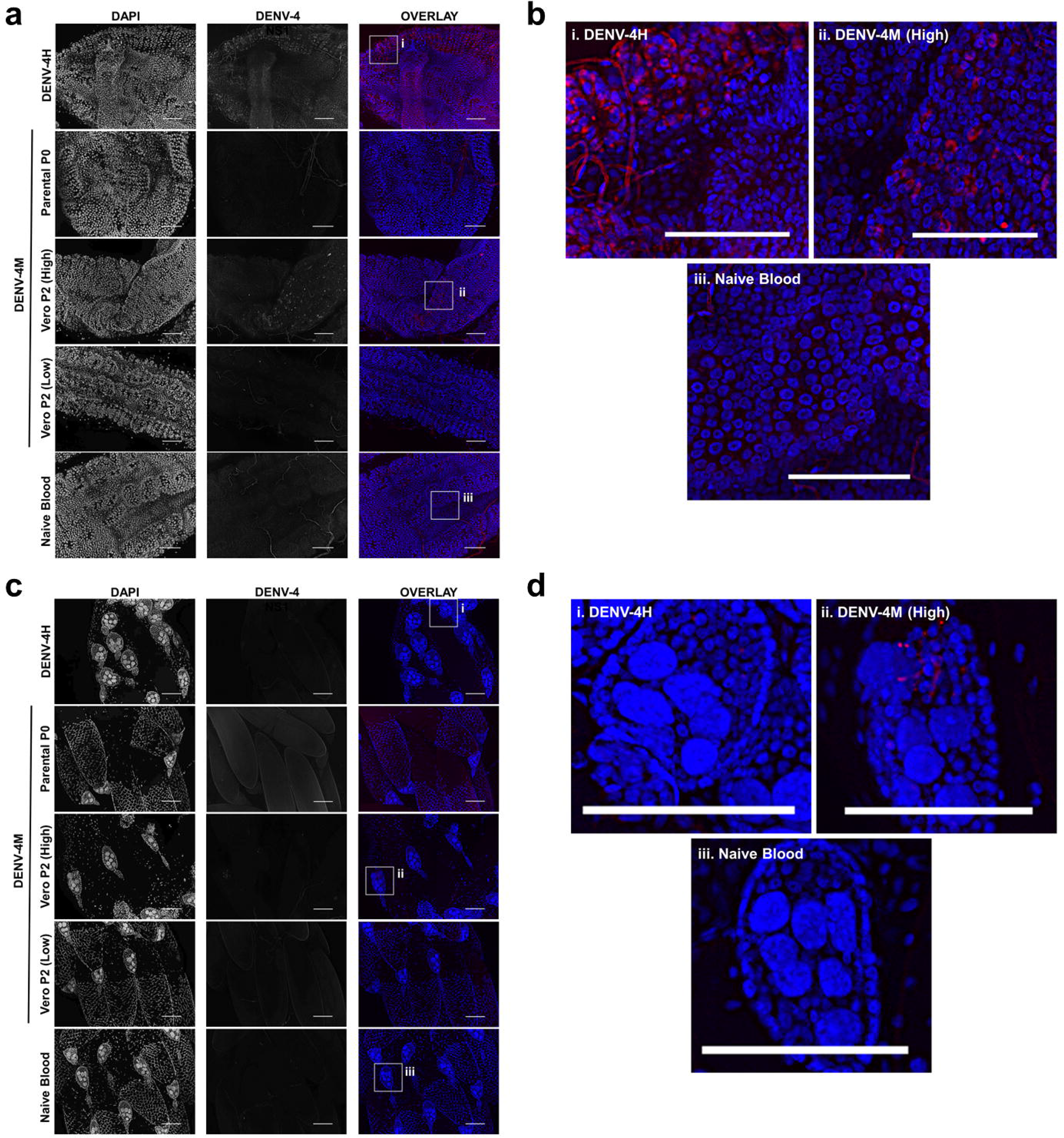
Multiple blood feedings do not increase DENV-4M infection prevalence or intensity. ORL mosquitoes infected with DENV-4M Vero E6 P2 were offered a second, uninfected blood meal at 4dpi (2BF) or not (1BF). At 14dpi, whole individual mosquitoes were collected for viral genome detection by rt-qPCR. Only individuals that were positive in both rt-qPCR technical duplicates were reported as positive. The proportion of positive mosquitoes to all mosquitoes tested is above each condition on the graph.

