## Supplementary Figures/Tables for "Ecology and transmission of a dengue virus serotype 4 identified in wild *Aedes aegypti* in Florida"

### 1 Supplementary Figures

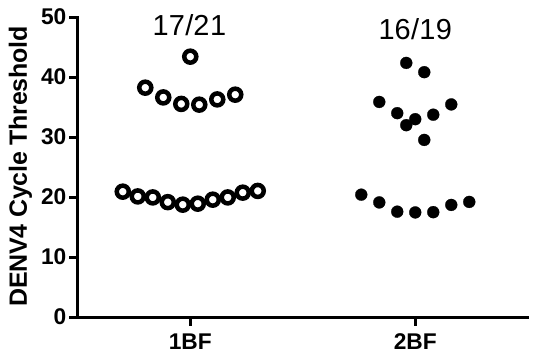

**Supplementary Figure 1.** **Multiple blood feedings do not increase DENV-4M infection prevalence or intensity.** ORL mosquitoes infected with DENV-4M Vero E6 P2 were offered a second, uninfected blood meal at 4dpi (2BF) or not (1BF). At 14dpi, whole individual mosquitoes were collected for viral genome detection by rt-qPCR. Only individuals that were positive in both rt-qPCR technical duplicates were reported as positive. The proportion of positive mosquitoes to all mosquitoes tested is above each condition on the graph.

### 2 Supplementary Tables

**Supplementary Table 1. Description of vertical transmission progeny pools.** Adult female F1 progeny of Vero E6 P2 DENV-4M infected females were pooled by rearing container, in pools of up to 25 individuals.

| **Pool #** | **1** | **2** | **3** | **4** | **5** | **6** | **7** | **8** | **9** | **10** | **11** | **12** |
| --- | --- | --- | --- | --- | --- | --- | --- | --- | --- | --- | --- | --- |
| **Replicate** | 1 | 1 | 1 | 1 | 1 | 1 | 1 | 2 | 2 | 2 | 2 | 2 |
| **# Females** | 4 | 25 | 22 | 16 | 25 | 3 | 23 | 4 | 2 | 11 | 14 | 1 |
| **DENV-4 positive** | No | No | No | No | No | No | No | No | No | Yes | No | No |

Supplementary Table 2: Sequences of primers used in construction and sequencing of the DENV-4M infectious clone.

| **Name** | **Sequence (5′ to 3′)** | **Note** |
| --- | --- | --- |
| DV4-570V | ATGTGTGAGGACACTGTC | For sequencing |
| DV4-1154V | AAGATGTCCAACGCAAGG | For sequencing |
| DV4-2323V | AGTGTTGTGGATTGGCAC | For sequencing |
| DV4-2911V | TCACGACCAACATATGGATG | For sequencing |
| DV4-3512V | TCTGTTGTGCCTGACCTTG | For sequencing |
| DV4-4204V | ATGATGTCCCTTTAGCAGG | For sequencing |
| DV4-4801V | AGACGTTCAGGTCCTCG | For sequencing |
| DV4-5384V | TCACCGATCCTTCCAGTG | For sequencing |
| DV4-5922V | ACAAGAAGACGACCAATAC | For sequencing |
| DV4-6550V | TAGCCTTACTAGGTGCTATG | For sequencing |
| DV4-7122V | GTGAACCCAACAACCTTG | For sequencing |
| DV4-7717V | AGCATGCAGTGTCTAGAG | For sequencing |
| DV4-8801V | TCAGGAAGAACAGGGATGG | For sequencing |
| DV4-9404V | CATGGAAGTTCAGCTCATC | For sequencing |
| DV4-9998V | AGTGTGGAACAGAGTGTG | For sequencing |
| DV4-10394V | GAAGCTGTACGCGTGG | For sequencing |
| DV4-F1 | attatacgaagttatattcgatgcggccgctaatacgactcac | Forward primer PCR of F1 |
| DV4-F2 | AGGAGCTATGCATTCAGCCC | Forward primer PCR of F2 |
| DV4-F3 | GACTTACACCCCGGAGCT | Forward primer PCR of F3 |
| DV4-F4 | AGcGTCGGGAAACATCGTGAG | Forward primer PCR of F4 |
| DV4-R1 | AGCGAGGGCTGAATGCATAG | Reverse primer PCR of F1 |
| DV4-R2 | CAGCTCCGGGGTGTAAGTC | Reverse primer PCR of F2 |
| DV4-R3 | AGAGCTCACGATGTTTCCCG | Reverse primer PCR of F3 |
| DV4-R4 | gtcgactctagaggatcccac | Reverse primer PCR of F4 |

Supplementary Table 3. Primer sequences for detection of dengue virus serotype 4 (DENV-4) (as per Santiago et al., 2013, doi:10.1371/journal.pntd.0002311).

| **Description** | **Name** | **Sequence** | **Fluor/Quencher** | **Position** | **Product size (bp)** | **Target** |
| --- | --- | --- | --- | --- | --- | --- |
| DENV-4 forward | D4-F_CDC | TTGTCCTAATGATGCTRGTCG |  | 884-904 | 89 | DENV4 prM |
| DENV-4 reverse | D4-R_CDC | TCCACCYGAGACTCCTTCCA |  | 953-973 |  |  |
| DENV-4 probe | D4-Pr_CDC | TYCCTACYCCTACGCATCGCATTCCG | FAM/BHQ-1 | 939-965 |  |  |
